# The Proximity Prediction Hypothesis: How predictive coding of CT-touch explains Autonomous Sensory Meridian Response and its therapeutic applications

**DOI:** 10.1101/2025.09.02.673729

**Authors:** Josephine R Flockton, Cade McCall, Catherine E. J. Preston

## Abstract

Autonomous Sensory Meridian Response (ASMR) is a pleasant tingling sensation felt across the scalp and neck, widely reported to reduce anxiety and improve sleep. The Proximity Prediction Hypothesis (PPH) is the first comprehensive predictive coding model explaining ASMR’s underlying neural mechanism. PPH posits that near-field acoustic cues from common ASMR triggers (e.g., brushing sounds, whispered speech) engage the audio-tactile Peripersonal Space Network, generating a top-down prediction of gentle C-tactile (CT) touch on CT fibre-rich skin of the scalp and neck. This prediction suppresses locus coeruleus (LC) arousal and increases vagal output, offering a mechanistic explanation for the phenomenon’s therapeutic benefits. In a subjective-experience survey (N = 64), ASMR-labelled trials were rated significantly more pleasant but only slightly more arousing than controls. Pleasantness predicted both the presence and intensity of tingles, supporting PPH’s core claim that hedonic value, rather than sympathetic activation, drives the graded somatosensory response. PPH situates ASMR within the Neurovisceral Integration framework, predicting measurable Central Nervous System-Autonomic Nervous System (CNS-ANS) markers (beta-band desynchronisation in the posterior insula and proportional increases in high-frequency heart rate variability with tingle intensity). It further predicts reduced LC activity during ASMR, stronger effects in individuals with high interoceptive prediction error (e.g., anxiety, autism), and attenuation of tingles when spatial proximity cues are removed. By integrating auditory proximity, CT-touch anticipation, and autonomic regulation into a single predictive-coding account, PPH provides a unified, testable framework for explaining ASMR, offering a blueprint for translating this sensory phenomenon into targeted, evidence-based interventions for anxiety and sleep disorders.

## 1. Introduction

Autonomous Sensory Meridian Response (ASMR) is a sensory phenomenon characterised by a pleasant tingling sensation felt across the scalp and often moving down the back of the neck, elicited by very specific stimuli. The sensation is triggered by auditory and/or audiovisual cues. Sounds that induce ASMR are varied and broad ranging, but the most popular triggers are slow, whispered speech, rhythmic hair brushing and tapping sounds (Barratt and Davis, 2015; Fredborg et al., 2021; Poerio et al., 2018). Alongside this, ASMR is also elicited via videos on media sharing platforms like YouTube where content creators use objects or their own voices to produce sounds that trigger the response in listeners. This is done by placing the camera and microphone near to the performers’ mouths or hands while they whisper or manipulate objects to make noises into the microphone. Over the past decade, ASMR content has transitioned from a niche phenomenon to a mainstream YouTube staple. As of 2022, there were approximately 500,000 ASMR-focused channels and an estimated 25 million ASMR videos on the platform, illustrating the breadth and scale of its cultural reach. Many ASMR videos fall into role-play genres that simulate close personal attention, including hairdresser visits, spa treatments, makeup application, doctor’s appointments, and other interpersonal care scenarios.

This popularity is seemingly driven by perceived benefits from experiencing the ASMR phenomenon, which go beyond the initial pleasant sensation. Survey work with hundreds of viewers found that 98% reported using ASMR for relaxation, 82% to help fall asleep, and about 70% to reduce stress or anxiety (Barratt and Davis, 2015, N=475). In laboratory follow-ups, participants who experience tingles report lower state-anxiety scores and improved mood up to thirty minutes after listening (Fredborg et al., 2021) suggesting the phenomenon provides more than just a pleasant distraction during the tingling experience itself and offers longer term affective benefits to those who enjoy it. These self-reports have also been scaffolded by physiological evidence. In a within subjects study that compared tingling to non-tingling segments of the same videos, Poerio and colleagues (2018) found a reliable heart rate deceleration accompanied by an increase in high frequency heart rate variability (HF-HRV). HF-HRV is a widely accepted non-invasive index of parasympathetic nervous system activity, often associated with states of calm and relaxation. Specifically, greater HF-HRV reflects increased vagal influence on the heart, indicating a shift toward physiological rest and recovery.

Recent work by Hozaki and colleagues (2025) extends this evidence using finger photoplethysmography (PPG), which not only captures pulse rate but also pulse wave amplitude, a measure of peripheral blood flow and vascular tone. In their study, both ASMR and nature videos reduced pulse rate relative to baseline, but ASMR produced significantly greater reductions. Moreover, ASMR was associated with increased pulse wave amplitude, consistent with peripheral vasodilation. Because vasodilation reflects parasympathetic dominance over vascular tone, these PPG findings complement HR and HRV evidence by demonstrating that ASMR’s autonomic effects extend beyond cardiac regulation to include vascular relaxation, reinforcing the interpretation of ASMR as inducing a coordinated parasympathetic shift. These parasympathetic-shift indicators are consistent with reduced sympathetic outflow, potentially reflecting suppression of tonic LC-noradrenaline activity. While PPG cannot directly index LC firing, the observed combination of bradycardia and peripheral vasodilation aligns with the physiological profile expected when LC tone is downregulated in contexts of perceived safety and calm.

Together, LC and vagus activity provide a functional gauge of the autonomic balance, where increased HF-HRV signals stronger vagal (parasympathetic) influence and PPG indices, specifically reduced pulse rate and increased pulse wave amplitude, indicate reduced sympathetic vascular tone. The fact that both cardiac (HF-HRV) and vascular (PPG) measures tilt in the same parasympathetic direction during ASMR grounds the present hypothesis in an established brain-body axis.

Although most research on ASMR has focused on mood benefits, some survey studies have revealed that sleep improvement is also a strong motivation for listening in many people. In Barratt and Davis’s (2015) 475 participant survey, 82% of responders reported using ASMR videos ‘often’ or ‘always’ to fall asleep faster. A later large-scale online study (Smejka and Wiggs, 2021; N = 1,037) found that ASMR viewing improved relaxation and mood across participants who did and did not suffer from insomnia. Although improvements were strongest in those who experienced tingles, no significant differences emerged between insomniacs and other groups in their response magnitude.

A mechanistic account is needed to link three disparate elements of the ASMR phenomenon: the acoustic character of the triggers, the subjective percept of pleasant scalp tingles, and the body-wide calming represented by physiological correlates like HRV and PPG, as well as reported mood and sleep benefits. A natural starting point is the Neurovisceral Integration (NVI) Model (Thayer and Lane, 2000). NVI frames mental state regulation as a dialogue between the cortical central-autonomic network (CAN) and subcortical autonomic nuclei. When the dialogue is smooth, indexed by high vagal tone and HF-HRV, the organism is flexible and resilient; when it is disrupted, anxiety and rumination flourish. Within this hierarchy the locus coeruleus functions as a noradrenergic ‘gain knob’; meaning elevated tonic LC firing biases the body toward sympathetic readiness, whereas a drop in LC tone likely lifts inhibition over the dorsal-motor nucleus of the vagus (DMV) and permits parasympathetic dominance, and calm. In a way, the LC and the vagus operate a seesaw-like regulatory axis that modulates perception and bodily state between arousal and relaxation. Existing ASMR findings, such as HRV increase during tingling and subjective experience reports, fit this framework, implying vagal activation and a downshift in LC tone. Yet no published stepwise neural model currently explains how auditory stimuli like whispers or brushing sounds could initiate this regulatory shift, let alone generate a tingling sensation across the scalp as a consequence.

Despite the range of auditory triggers that can elicit ASMR in listeners, one property which they arguably all have in common is that they can be categorised as proximal, near-ear stimuli, rich in spatial cues, illustrated by three key acoustic features shown across the literature. First, very large interaural level differences (ILDs) and sub-millisecond interaural time differences (ITDs) signal that the sound source is only a few centimetres from the listener’s head. ASMR YouTube video recordings are typically made with binaural ‘dummy-head’ microphones whose fake pinnae and ear canals preserve these cues; playback over loudspeakers reduces them, but headphones, through which 90% of listeners choose to experience ASMR (Barratt and Davis, 2015; N = 475), deliver them unchanged, recreating the illusion that a hand or brush is at the ear. Second, the spectrum is colour-shifted by head-shadowing, meaning high frequencies above 8 kHz roll off steeply in the contralateral ear, a cue which listeners tend to interpret as indicating close spatial proximity (Begault,1994). Third, ASMR content creators often favour slow amplitude envelopes and low overall sound pressure levels. This means that the loudness of the signal rises and falls gradually, over hundreds of milliseconds or more, rather than in sharp, percussive bursts. A whispered phrase, a brush stroke across a microphone, or a series of soft taps typically shows a smooth, rounded waveform without abrupt transients. In addition, keeping the overall sound pressure level low ensures the audio remains intimate and non-startling, helping listeners maintain a relaxed, parasympathetic state; louder levels would recruit the middle ear reflexes and risk activating the sympathetic ‘alerting’ system, which would contradict the calming goal of ASMR.

These findings converge on an interesting idea, that ASMR stimuli may work to convince the auditory system that an object is virtually approaching or touching the ear or scalp, in the absence of any real physical contact. A mechanistic model must therefore account for the special spatial signature of these sounds, then explain how such proximity information could cascade into both the tingling percept and the parasympathetic shift measured in HRV and through reported improvements in mood and sleep. This paper proposes a Proximity Prediction Hypothesis (PPH) to integrate the audio-tactile features mentioned above, with the NVI framework, arguing that near-field sounds pre-activate the brain’s Peripersonal Space Network and prompt a top-down prediction of impending gentle CT-touch on the scalp.

Valtakari et al. (2019) observed that ASMR experiences are accompanied by pupil dilation, while Poerio et al. (2018) reported increased skin conductance responses (SCR) during tingling segments compared to control periods. Both pupil dilation and SCR are well-established markers of sympathetic nervous system activity, indicating that ASMR is not a purely parasympathetic phenomenon. This has caused some debate in the literature, given its reportedly calming profile. However, as McGeoch and Rouw (2020) note, the combination of heart rate deceleration and increased SCR suggests both sympathetic and parasympathetic involvement and, because eccrine sweat glands (underlying SCR) receive only sympathetic innervation, while the heart is dually innervated by both sympathetic and parasympathetic pathways, the net decrease in heart rate points to an overall shift toward increased vagal tone. This aligns with the PPH model, in which pupil dynamics in ASMR reflect a transition from orienting to affiliative calm. Near-ear cues initially engage the LC-noradrenaline system, producing a transient pupil dilation to enhance sensory gain. As peripersonal space and CT-afferent touch predictions converge, tonic LC activity is suppressed and parasympathetic output dominates, leading to heart rate deceleration, increased HF-HRV, feelings of calm, and we predict, eventual pupil constriction. This biphasic pattern accommodates both sympathetic (early attentional) and parasympathetic (later calming) components, supporting the interpretation of ASMR as a flow state (Peifer et al., 2014) of ‘relaxed alertness’ characteristic of safe, affiliative proximity.

After explaining the theoretical background, current evidence in the area will be collated and assessed in the context of the PPH model. Then, we report original illustrative survey data from sixty-four listeners in an immersive ASMR listening study, demonstrating that hedonic valence drives the tingling experience and its intensity, thus providing empirical support for the PPH model. Clinical applications and the reported benefits to mental health and sleep in ASMR experiencers will be discussed with the PPH model and CNS-ANS integration in mind. Future research will be suggested to test the theory, with falsifiable predictions for findings across CNS-ANS research, encompassing heart rate variability, pupil-indexed LC dynamics, and beta band neural signatures, in behavioural, EEG, and MEG studies, if the model is to be supported.

## 2. Theoretical Foundations

### 2.1 The interoceptive brain and predictive coding

According to Interoceptive Predictive Coding accounts (Critchley and Harrison, 2013; Barrett and Simmons, 2015), cortical areas generate continuous, probabilistic forecasts (or ‘priors’) about what the viscera, skin, and muscles should feel like in the next few hundred milliseconds. Incoming afferent data are compared with these priors and any difference found is the prediction error signal (Feldman and Friston, 2010). A close match is desired; a mismatch registers as physiological surprise and, when this is chronic, can lead to anxiety. When the incoming signal and priors match (or the error is negligible), this implies that the sensory world is unfolding as expected. Most of this comparison takes place in areas such as the posterior and anterior insula, which influence autonomic nuclei in the brainstem. The posterior insula receives raw interoceptive input, constructs a sensory map of the body, and forwards that map to the anterior insula, where predictions and errors are integrated with the affective context (Critchley and Harrison, 2013). When the match between the prior and signal is close, and the prediction error is small to negligible, for example, if you feel the gentle pressure that you expected while holding a cup in your hand, the anterior insula sends an inhibitory signal to the locus coeruleus (LC), the brainstem hub for noradrenaline release. In simple terms, this inhibits the LC’s usual role in promoting arousal and vigilance. As tonic LC firing drops, its noradrenergic brake on the dorsal-motor nucleus of the vagus (DMV) is lifted. The result is increased vagal output and a rise in high frequency heart rate variability (HF-HRV), the parasympathetic signature of calm suited for rest, digestion, and affective ease (Samuels and Szabadi, 2008).

This low precision gate explains everyday illusions like in the phantom phone buzzing phenomenon, where people report feeling their phone vibrate even when it is not; a strong learned prior (‘my phone is about to vibrate’) meets either minimal somatic noise or no detectable cutaneous input at all. Because any residual error is labelled as low precision, the posterior insula fills in the expected buzz with a somatosensory echo, a phantom vibration, the anterior insula reports ‘prediction fulfilled,’ and the LC-vagus axis remains calm (Lin et al., 2013). Virtual reality touch has a similar mechanism where viewing a virtual stick stroking a forearm that you associate with your own body in virtual reality produces tingles in 89% of users despite zero skin input on their actual arm in real life, because the visual prior overwhelms the ill-defined cutaneous error (Pilaciński et al., 2023), it is more likely that you are being touched and it is light and not hugely noticeable, than that all other, more reliable, priors are wrong in anticipating that touch when your previous experience and the visual input suggests it is very likely. In both cases of touch illusions, the visual or contextual prior overwhelms the ambiguous tactile input. The cue is interpreted as consistent with expected gentle touch but not clear enough to generate high precision error, allowing the prior to dominate. Touch is considered ill-defined in these circumstances because the sensory evidence is either absent, ambiguous, or delivered through a channel (e.g. auditory or visual) that does not strongly engage tactile precision mechanisms. When this occurs, the brain is more likely to accept the predicted sensation and resolve the ambiguity in favour of the expected state.

Crucially, ‘precision’, the brain’s estimate of sensory reliability i.e. its confidence in the fidelity of a particular sensory channel, modulates how much any given error matters. High precision channels (e.g. retinal contrast, a pin-prick sensation) deliver errors that are hard to ignore; low precision channels however (faint rustling, diffuse light pressure) deliver errors that can be treated as background noise. Here we suggest that, when the brain issues a strong top-down prior like ‘I am about to feel a gentle stroke’ and the incoming signal is fuzzy, delayed, or absent, the mismatch is labelled as low precision. In that case the posterior insula may simply fill in the expected sensation itself and send a ‘prediction fulfilled’ message upstream. Because the error never gains salience, the anterior insula does not escalate to the LC, tonic LC firing falls, and the vagal brake is released even though no physical touch ever occurred. This series of predictive, neurophysiological events, from sensory prior to vagal activation, forms the basis of the Proximity Prediction Hypothesis (PPH) cascade, a stepwise model proposed to explain how the characteristic calm and tingling response of ASMR can arise from purely auditory cues. Each element of this cascade is explored in subsequent theoretical sections and visualised in Figure 1 below.

**Figure 1.**
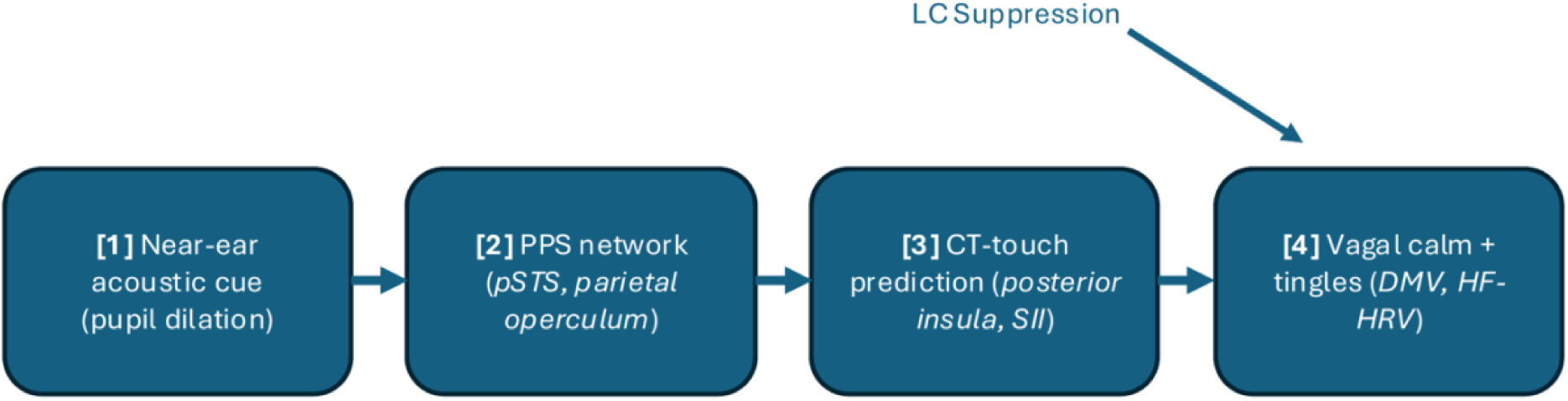
Proposed cascade of the Proximity Prediction Hypothesis (PPH). A near-ear sound activates peripersonal space (PPS) networks, which forwards a C-tactile (CT) touch prediction to somatosensory and interoceptive regions; confirmation of that prediction suppresses locus coeruleus (LC) tone, disinhibits the vagal system, and generates parasympathetic calm and tingling sensations. **[1] Near-ear acoustic cue → PPS detection:** binaural whispers and brushing sounds carry strong interaural time and level differences, interpreted by the posterior superior temporal sulcus (pSTS) and adjacent areas as proximal, human-origin sounds (Schürmann et al., 2006; Belin et al., 2000; Warren and Griffiths, 2003); at this early orienting stage, sympathetic attentional mechanisms such as pupil dilation are transiently recruited to enhance sensory gain (Valtakari et al., 2020). **[2] PPS network → CT-touch prediction:** pSTS and parietal operculum project to the posterior insula and secondary somatosensory cortex (SII), simulating tactile consequences of perceived social proximity, especially on CT-rich scalp/neck regions (Löken et al., 2009; Gazzola and Keysers, 2009). **[3] Accurate prediction → LC suppression and vagal disinhibition:** minimized prediction error reduces anterior insula drive to the LC, lowering tonic noradrenaline and lifting inhibitory control over the dorsal motor nucleus of the vagus (DMV), increasing parasympathetic tone and yielding cardiac deceleration and increased high-frequency HRV (Paulus and Stein, 2006; Samuels and Szabadi). **[4] Conscious correlate; the tingles:** pre-activation of insula/SII yields a synchronous, spatially diffuse cortical volley experienced as a tingling somatosensory echo of predicted contact.

A similar process is proposed more generally in the Somatic Error Hypothesis (Van den Bergh et al., 2017), where the brain reduces prediction error by generating bodily sensations that match an expected state. While this mechanism is typically invoked to explain chronic symptoms in somatising disorders, here we extend its logic to a benign interoceptive illusion felt by those who experience ASMR.

### 2.2 The audio-tactile fabric of peripersonal space

Prediction in this case does not operate in isolation, it is shaped by multisensory maps of Peripersonal Space (PPS), which can be thought of as a bubble of 20-30 cm space surrounding the body where approaching objects are most likely to make contact.

Importantly, PPS is not a simple distance gradient; it behaves like a biological boundary. Stimuli presented just inside the bubble elicit abrupt neural and behavioural changes, whereas equally small decrements in distance once the stimulus is outside the bubble have little effect (Làdavas and Serino, 2008; Serino, 2015). A substantial body of multisensory work shows that the brain treats a near-ear sound as a potential touch event.

Early single-unit electrophysiology in macaque monkeys revealed a class of multisensory neurons in ventral premotor and parietal regions, including the ventral intraparietal area (VIP), that integrate tactile, visual, and auditory signals relevant to peripersonal space.

Auditory cues alone can activate neurons in peripersonal space-sensitive regions, including the VIP, for instance, Graziano, Reiss, and Gross (1999) reported that broadband noise sources moving toward the head, from 70 cm to 10 cm, caused multisensory neurons in VIP to fire more vigorously than when those same stimuli moved within far space. This indicates that approaching sounds, even in the absence of visual input, can signal potential contact and recruit defensive spatial coding. Moreover, Avillac and colleagues (2007) demonstrated that VIP neurons integrate visual and tactile input when sensory events are spatially and temporally aligned. These neurons often integrate tactile and auditory information, reinforcing the idea that auditory proximity cues are biologically relevant indicators of incoming contact.

Importantly, VIP neurons respond to stimuli that occur both on the body (i.e., within a neuron’s tactile receptive field) and just beyond it, typically within a few tens of centimetres. This alignment of visual and somatosensory receptive fields reflects a body-centred coding of nearby space, a neural basis for anticipating contact (Colby et al., 1993; Rizzolatti et al., 1997).

Magnetoencephalography supports the idea that ASMR-like stimuli can activate such somatosensory regions. Schürmann et al. (2006) played realistic sounds resembling haircut and water-dripping scenarios and found beta-band desynchronisation in the secondary somatosensory cortex (S2). This effect is supported across broader studies. Canzoneri and colleagues (2012) found that sounds approaching the hand significantly accelerated tactile responses once perceived within peripersonal space. A meta-analysis by Holmes et al. (2020) confirmed a modest (15 ms) reduction in tactile reaction times when sounds occurred near the body versus farther away, although they noted variability and small effect sizes.

Additionally, Taffou and colleagues (2014) demonstrated that looming ‘rough’ sounds, those with threat-like acoustic properties, expanded the effective PPS boundary, triggering tactile facilitation at greater distances than smoother sounds. Together, these findings support the notion that sound proximity is a potent modulator of sensory integration and may help explain how ASMR content elicits embodied responses despite being purely auditory.

These findings demonstrate that the posterior STS, inferior parietal cortex, and the parietal operculum behave like proximity detectors, amplifying their response when an auditory object crosses the PPS boundary, and is therefore likely to make physical contact. This supports the notion that sound proximity is a potent modulator of sensory integration and may explain how ASMR content elicits embodied responses despite being purely auditory. Such proximity-sensitive firing is proposed to represent the first node in the PPH cascade, the moment when the brain interprets near-ear sounds as predictive of imminent affective touch, triggering downstream autonomic changes detailed in the next sections (see Figure 1).

### 2.3 C-tactile afferents and the mechanism behind affective touch

If, as PPH suggests, ASMR is occurring through prediction of affective touch, it is important to consider what exactly the brain is predicting and how that links to the reported ASMR experience. When contact does occur on the skin, it is detected by at least two tactile channels. Fast, myelinated A-β fibres handle discriminative features, conveying facts about the touch, like location, texture, and force, whereas C-tactile (CT) afferents are slow, unmyelinated fibres that overwhelmingly tend to innervate hairy skin regions.

Microneurography shows that CT afferents respond optimally to gentle stroking at 1-10 cm s⁻¹, with a firing peak at around 3 cm s⁻¹, which is exactly the velocity of social grooming strokes in primates (Löken et al., 2009; Ackerley et al., 2014). Their firing rate predicts subjective pleasantness and drives oxytocin release, posterior-insula activation and a parasympathetic drop in heart rate (Ackerley et al., 2014; Pawling et al., 2017).

Human CT afferents have been recorded in scalp, face, forearm, abdomen and thigh areas (McGlone et al., 2014) and show a clear preference for hairy skin. While detailed follicle density maps are scarce, regions such as the scalp midline, nape, and upper back are widely associated with social grooming in primates and are plausible candidates for dense CT innervation (McGlone et al., 2014). These zones are therefore likely to be particularly well populated by CT-touch fibres. They are also prime cortical targets for affective touch, where the brain predicts a gentle, grooming-like sensation to land. Crucially, this is indeed where the ASMR tingling sensation is reported to be localised: the scalp, face, neck, and upper back (Barratt and Davis, 2015; Poerio et al., 2018; Lochte et al., 2018).

Behavioural data echo the physiology; in barbary macaques, bouts of allogrooming (a prosocial behaviour where animals of the same species groom one another) lower basal cortisol and heart rate within minutes (Shutt et al., 2007). In humans, five minutes of scalp massage at CT-optimal velocity produces a significant HF-HRV increase and self-reported anxiety reduction in Spielberger state-anxiety test scores (Diego and Field, 2004).

Consistent with this, a 45-minute relaxation massage before bed has been shown to enhance sleep efficiency in individuals with insomnia (Ntoumas et al., 2025). Moreover, meta-analytic evidence indicates that interventions involving head touch specifically, such as face or scalp massage, may confer particularly strong physical and mental health benefits (Packheiser et al., 2024), reinforcing the potential relevance of affective touch to ASMR-related somatosensory modulation. As the second stage of the cascade, CT-touch predictions anchor the brain’s expectation of safety and interpersonal care. Taken together, these findings establish CT-touch as a hedonic, anxiolytic, and sleep-promoting modality, and identify the scalp and neck as privileged substrates for such contact, exactly the locations where ASMR listeners report feeling their tingles.

One complementary, structural account of ASMR has been offered by McGeoch and Rouw (2020), who propose that ASMR may involve synesthetic cross-activation between the primary auditory cortex (A1) and affective-touch maps in the dorsal posterior insula (dpIns). This pathway could provide the substrate for the CT-touch predictions outlined in the PPH: under proximal, interpersonal conditions, near-ear sounds may recruit such regional cross-activation to simulate gentle social touch, triggering posterior-insula activity and vagal engagement. Unlike the PPH, however, their model does not address the state-dependent gating, peripersonal space integration, or temporal autonomic cascade that determine *when* and *how* this cross-activation occurs. The two perspectives can therefore be viewed as complementary, with cross-activation describing the same plausible neural route and the PPH specifying the predictive coding logic and dynamic conditions under which that route is engaged in the ASMR phenomenon. Furthermore, while McGeoch and Rouw’s hypothesis links auditory input to affective touch areas, it does not specify the computational mechanism by which tingles emerge, nor how such activation alone would produce the distinct, wave-like somatosensory echo characteristic of ASMR. The PPH extends this by proposing the predictive coding process and time-resolved neural signature capable of transforming such cross-activation into the tingling percept itself.

### 2.4 The social neurocognitive context of ASMR

If ASMR indeed reflects a prediction of affiliative touch, then understanding the social and cognitive conditions that shape those priors becomes crucial. ASMR triggers overwhelmingly reflect socially salient acts like whispering, soft-spoken instruction, and gentle, attentive behaviours, many of which imply close interpersonal proximity. These cues may be sufficient to evoke predictions of touch-like feedback, particularly in individuals predisposed to interpret such signals as affiliative or comforting.

Recent empirical studies have identified five principal ASMR trigger categories, all of which share a perceptual association with human interaction: (1) viewing individuals interact with objects, (2) watching socially intimate acts, (3) hearing soft repetitive sounds, (4) simulated social interaction, and (5) whispering or chewing (Smith et al., 2020; Fredborg et al., 2017). Even seemingly nonsocial triggers, such as tapping or crinkling, often co-occur with goal-directed behaviours that implicitly suggest a human source (Janik McErlean and Banissy, 2017). This convergence supports the idea that ASMR is scaffolded by social perceptual priors, often concerning caregiving or affiliative intent.

Recent work by Poerio and colleagues (2023) developed the ASMR Trigger Checklist (ATC), a validated tool for systematically identifying and categorizing common ASMR triggers, to assess how individuals respond to a wide range of sounds. They found considerable variability in which triggers reliably induced tingles across participants. Critically, the most potent triggers tended to be those that implied gentle, interpersonal interaction or close physical proximity, such as whispering or soft tapping. This variability is consistent with the precision-weighting mechanism proposed by the PPH; individuals may assign higher predictive value to particular sensory cues based on their internal priors about social intent, interpersonal closeness, or expected affective touch. The ATC therefore offers a structured way to assess which auditory signals carry predictive weight in different individuals, and why the same cue may trigger ASMR in one person but not another. These findings reinforce the notion that ASMR emerges from a socially grounded predictive model shaped by prior experience, attachment tendencies, and interoceptive sensitivity.

Most significantly for the PPH, Poerio’s study (2023) also showed that physical touch itself, rather than sound or visual cues, was not only the most commonly endorsed ASMR trigger reported to elicit a tingling sensation in participants (98%) but also the most intense, with minimal variation across individuals; the ATC subset of tactile and interpersonal triggers gave examples like ‘close-up movements directed at you’ and ‘light touch on your face e.g. make-up application’. This highlights that touch itself, whether anticipated or actively experienced is a core trigger for the ASMR tingling sensation, making the idea of a somatosensory echo even more plausible as it is clear that the tingling sensation is a ground truth for the phenomenon, not an abstract, novel response the brain is predicting. This emphasizes that the tingles are less of a bodily illusion, as some may argue, and more of a plausible sensory prediction based on what it does actually feel like when people are really being touched.

Importantly, Poerio and colleagues argue that online ASMR content should be seen as a simulation of real-world interpersonal encounters rather than as distinct from them, and that trait ASMR may be meaningfully defined by a person’s sensitivity to touch-related triggers. In this way, their work empirically supports the idea that ASMR operates through predictive interoceptive mechanisms shaped by tactile expectation and affiliative social context.

Consistent with this view, Gillmeister et al. (2022) demonstrated that gentle social touch enhanced the intensity and pleasantness of ASMR responses, but only in ASMR-experiencers, reinforcing the role of trait-dependent priors for affiliative interaction in driving the ASMR response. This final phase of the cascade, the culmination of proximity, touch prediction, and arousal regulation, is therefore likely shaped by an individual’s social priors, attachment style, and interoceptive sensitivity.

Neuroimaging studies further reinforce the social grounding of ASMR. Lee et al. (2020) found that during ASMR experiences, participants showed activation in brain regions implicated in social cognition and mental state simulation, including the posterior cingulate cortex, superior and middle temporal gyri, and the lingual gyrus. These areas are key components of the brain’s social mentalizing network, suggesting that ASMR may engage the same systems we use to interpret and internalize others’ intentions; particularly when those intentions are perceived as caring, attentive, or intimate.

Earlier work by Lochte (2018) proposed that ASMR may function as a vestigial grooming response, with Lochte going on to suggest that ASMR may be a polymorphic trait, a term used in evolutionary biology to describe a characteristic that is present in some individuals of a species but not all, due to genetic or developmental variability. Common examples include wisdom teeth or lactose tolerance, traits that were once adaptive, but are now only expressed in certain subsets of the population. If ASMR is indeed a polymorphic vestige of an ancestral grooming response, this could explain why only some individuals report experiencing tingles in response to specific stimuli. Again, aligning with the PPH model’s suggestion that ASMR emerges only when an individual’s internal predictive model assigns high precision to interpersonal proximity cues, a tendency that may itself vary across individuals based on neurocognitive, social, or interoceptive traits. These accounts offer an ethological framework for why ASMR stimuli elicit pleasure and calm in a specific subset of individuals.

That subset may be defined, in part, by individual differences in trait empathy and sensory-emotional inhibition. Janik McErlean and Banissy (2017) reported that ASMR experiencers tend to score higher on ‘Empathetic Concern’, suggesting a heightened sensitivity to social-affective cues. Others have found that ASMR is associated with reduced functional connectivity in the prefrontal cortex and default mode network (Smith et al., 2017; Fredborg et al., 2021), implying diminished top-down inhibition of incoming sensory-affective stimuli. In predictive coding terms, such individuals may assign greater precision to exteroceptive social cues while allowing these predictions to unfold with minimal suppression, creating fertile ground for ASMR to emerge.

## 3. The Role of Pleasantness in ASMR

If a near-ear whisper is effective because it forecasts a slow, pleasant interpersonal contact, then the strength of the ASMR response should depend on how rewarding that predicted contact feels, not on its sheer acoustic energy. Within the PPH framework, pleasantness (valence) is expected to determine two outcomes: whether a listener classifies a segment as ASMR at all, and how intense the tingles feel during the ASMR experience. This mirrors genuine affective touch, where C-tactile firing rates track subjective pleasantness (Löken et al., 2009) and hedonic ratings predict downstream effects on pain perception (Pawling et al., 2017). By analogy, ASMR tingles should scale with pleasantness because the posterior insula propagates stronger predictions of affective touch when the hedonic prior is stronger.

This valence-first logic is also illustrated by the content ecology of ASMR. The most watched videos on YouTube are spa, hairdresser, and make-up roleplays in which creators whisper reassurances, move brushes and scissors centimetres from the microphone, and enact a caretaking script. Such clips are maximising both near-field spatial cues and a social-grooming context, forming a strong hedonic prediction with minimal arousal load. Because the CT-touch prediction is intrinsically hedonic, a dominance of pleasantness over arousal would mirror the physiology of real affective touch. Microneurography shows that the firing rate of C-tactile afferents rises monotonically as stroking speed approaches the 3 cm s⁻¹ optimum and that subjective pleasantness ratings track this firing curve with an almost unit slope (Löken et al., 2009). Follow-up psychophysics demonstrated a similar scaling for behavioural impact; in Pawling et al. (2017) each one-point increase on a 10-point pleasantness scale produced an additional 0.9-point decrease in pain rating during concurrent heat stimulation, confirming that the more pleasant the predicted stroke, the stronger its sensory-affective consequence. In other words, CT-touch intensity is modulated by valence in exactly the way PPH predicts ASMR might be.

This account is further supported by recent behavioural data from Gillmeister et al. (2022), who found that ASMR responders reported significantly greater tingle intensity and pleasantness ratings in response to auditory ASMR triggers when accompanied by gentle interpersonal touch, whereas non-responders showed no such modulation. Notably, the strength of the tingle correlated more with pleasantness than with arousal, underscoring the centrality of hedonic predictions in driving ASMR’s intensity. Their findings align with the PPH in suggesting that social touch cues amplify ASMR not through generic arousal, but through affective reward mechanisms that may be trait-dependent.

Importantly, this does not mean that arousal is irrelevant to ASMR. Some triggers may increase both pleasantness and arousal, and the role of arousal remains equivocal. What PPH predicts, however, is that pleasantness will be the primary driver of whether a sound crosses the tingle threshold and of how strong those tingles become.

The next section tests this prediction directly, using trial-level behavioural data on pleasantness, arousal, and ASMR reports from an original survey dataset by the authors.

## 4. Illustrative Behavioural Evidence

### 4.1 Method

#### 4.1.1 Participants

Undergraduate students (N=64) from the University of York were recruited to participate in this study, there was no requirement for having any previous experience of ASMR, in order to reduce expectation bias about experiencing the phenomenon, without simply excluding those who may have listened before. An information sheet provided before the initial EEG study (in which each person had participated before completing this subsequent questionnaire study) explained the definition of ASMR to all participants from the start. The only exclusionary criteria was ensuring participants were over the age of 18, and that they had no known previous negative experience of ASMR sounds that would make them likely to feel discomfort when listening.

#### 4.1.2 Stimuli

Sound clips were used from a wider EEG study, these clips were produced by an ‘ASMRtist’ content creator who had given consent for the use of this selection of sounds that have been known to induce ASMR and are popular with their listening audience including paper folding, tapping, stroking sounds - no mouth sounds were used in order to avoid triggering misophonia in any listeners inadvertently. In the initial stimuli presentation study, each trial consisted of a 30 second sound stimulus, with a 5 second interstimulus interval (ISI), and all stimuli were randomly presented. Auditory stimuli were presented via Sennheiser HD280 Pro Dynamic Hi-Fi Stereo headphones connected to an AudioFile zero-latency triggerable audio stimulus generator (Cambridge Research Systems) during the experiment. The EEG experiment was conducted in a sound-attenuated room with participants lying in a supine position on a massage table to promote comfort and minimise movement. There were 18 sound stimuli (13 experimental sounds created by an ASMRtist and 5 control sounds of ambient traffic noise recorded by the experimenters) presented in a single randomised block during the original experiment. For the purpose of the presently presented questionnaire study, participants were asked to remember what their response to those experimental sounds in the original study had been, in a brief questionnaire after the EEG session had finished. Participants were prompted with shorter 5 second clips of the same experimental sounds in the questionnaire, that they listened to through the same headphones, in the same sound-attenuated room, to help them remember each clip when they were asked questions regarding each of those 13 experimental sounds.

#### 4.1.3 Procedure

After completing the EEG recording session in which they listened to 13 ASMRtist-created experimental sounds and the 5 control sounds mentioned above, participants were presented with a post-experience digital questionnaire assessing their subjective responses to the experimental stimuli. The questionnaire replayed the first 5 seconds of each of the 13 experimental sounds (presented in randomised order, just like during the original session) and asked participants to indicate, for each sound, whether they believed they experienced ASMR (“Yes” or “No”). The participants were informed of the definition of ASMR in the information sheet provided before the EEG experiment, and there was no requirement to have been familiar with ASMR or know if you could experience it, to sign up for the study or subsequent follow-up questionnaire. If “Yes” was selected to suggest ASMR had been experienced for any sound, participants were prompted to provide a retrospective estimate of tingle intensity on a 0-10 scale, if they could recall the sensation. All participants also used on-screen sliders to rate the pleasantness and arousal associated with each sound on continuous scales from −250 (extremely unpleasant or calming) to +250 (extremely pleasant or arousing) whether they experienced ASMR for that sound or not. The survey was completed immediately after the listening to the full clips, while participants remained in the sound-proof testing room environment, to minimise memory decay and distraction, and using the same headphones for sound clip delivery. Not all participants provided tingle intensity ratings for each sound, as this question was optional and conditional on an ASMR report as well as their memory of it.

#### 4.1.4 Statistical analysis

To investigate what drives whether a sound elicits ASMR, and the strength of the associated tingling sensation, mixed-effects regression models were implemented in R (version 4.4.0) using the lme4 (Bates et al., 2015), lmerTest (Kuznetsova et al., 2017), and broom.mixed (Bolker et al., 2024) packages. Predictors were z-scored to aid interpretation and comparability. A logistic mixed-effects model was used to predict ASMR classification (either Yes or No) from pleasantness and arousal ratings, with random intercepts for participant and sound. A subsequent model tested whether the effect of pleasantness on reported ASMR experience was moderated by arousal using an interaction term.

For trials where participants reported experiencing ASMR and rated its intensity, a linear mixed-effects model was used to predict tingle strength from pleasantness and arousal, again including an interaction term in a follow-up model. Visualizations were created using ggplot2 (Wickham, 2016) and ggeffects (Lüdecke, 2018), with predicted probability heatmaps and scatter plots depicting the effects of predictors across trials and sound clips.

### 4.2 Results

#### 4.2.1 What drives the ASMR decision?

A mixed-effects logistic regression model was fit with ASMR classification (Yes or No) as the outcome and z-scored pleasantness and arousal as fixed effects, with random intercepts for participant and sound.

Pleasantness emerged as a strong positive predictor of ASMR reports (β = 2.07 ± 0.23, *z* = 8.86, *p* < .001), while arousal showed a non-significant negative trend (β = –0.29 ± 0.17, *p* = .092). This suggests that hedonic valence, rather than activation level, primarily drives the ASMR decision.

An interaction term between pleasantness and arousal was also tested to assess whether arousal modulated the effect of pleasantness. However, the interaction was not statistically significant (β = 0.20 ± 0.15, *p* = .194), and did not improve model fit (likelihood ratio test: χ²(1) = 1.68, *p* = .195). Therefore, the probability of classifying a sound as ASMR was strongly driven by pleasantness across the full arousal range.

Figure 2 below visualizes the predicted probability of reporting ASMR as a function of z-scored pleasantness and arousal. The near-vertical gradient in predicted probabilities underscores the dominance of hedonic valence in the ASMR decision; increases in pleasantness robustly predict ASMR reports across the full arousal range, while arousal adds minimal predictive power.

**Figure 2.**
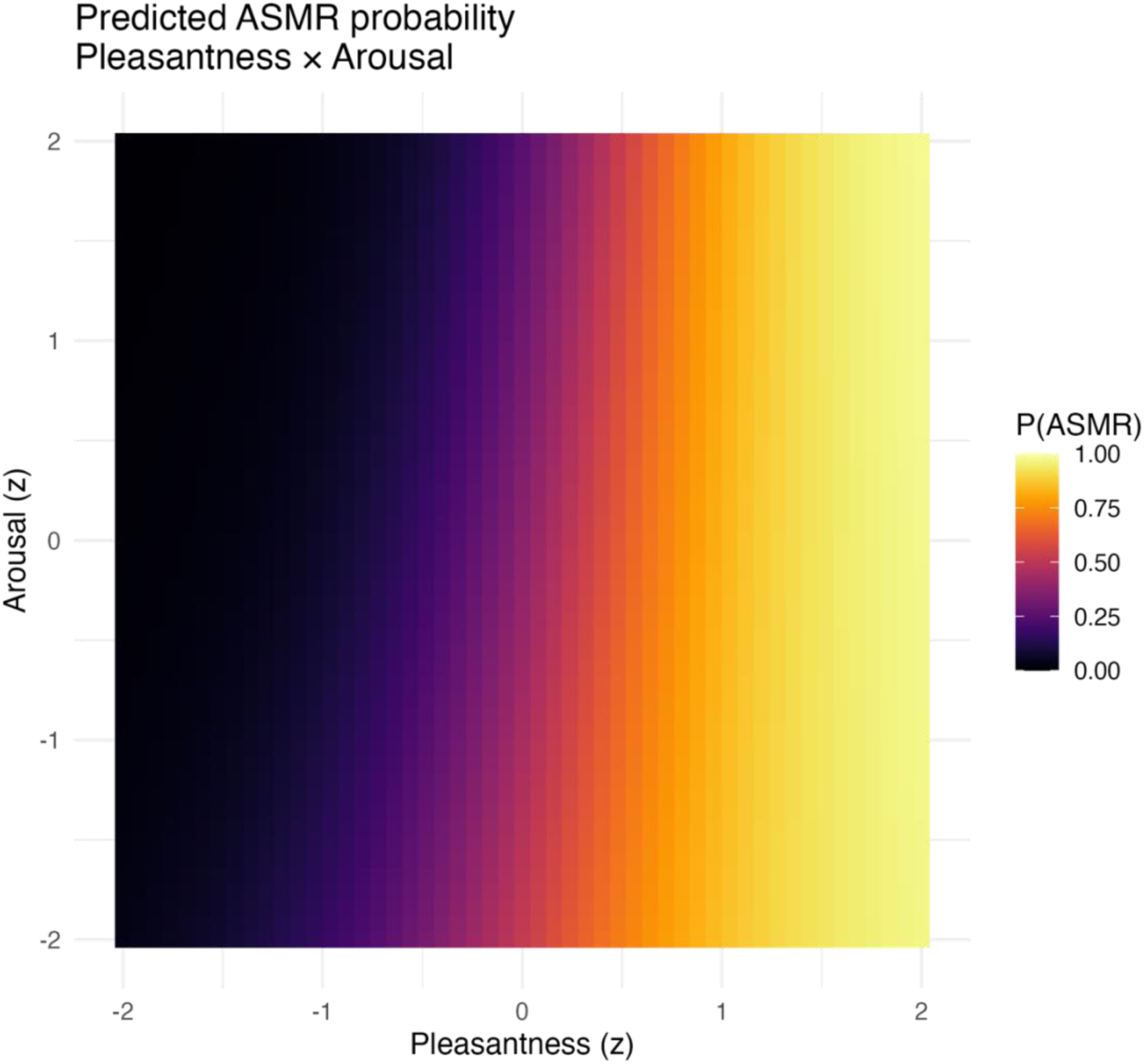
Predicted probability of ASMR classification as a function of z-scored pleasantness (x-axis) and arousal (y-axis). Colours show predicted probabilities from a logistic mixed-effects model with random intercepts for participant and sound (N = 64). Near- vertical contour lines indicate pleasantness as the dominant predictor, with minimal modulation by arousal.

Table 1 below presents the fixed effect estimates from the full logistic regression model, including the interaction term.

**Table 1.** Fixed-effect estimates from the logistic mixed-effects model predicting ASMR classification from z-scored pleasantness and arousal (including their interaction), with random intercepts for participant and sound.

| Term | Estimate | SE | Z | *p* | 95% CI (Low) | 95% CI (High) |
| --- | --- | --- | --- | --- | --- | --- |
| (Intercept) | -0.499 | 0.424 | -1.180 | 0.239 | -1.330 | 0.332 |
| Pleasantness (p_z) | 2.070 | 0.233 | 8.860 | 0.000 | 1.610 | 2.520 |
| Arousal (a_z) | -0.293 | 0.173 | -1.690 | 0.091 | -0.633 | 0.047 |
| Pleasantness x Arousal (p_a:a_z) | 0.196 | 0.151 | 1.300 | 0.194 | -0.100 | 0.492 |

#### 4.2.2 Tingle intensity

On 136 ASMR-positive trials with self-rated intensity scores, tingle strength increased linearly with pleasantness (β = 1.06 ± 0.22, t = 4.71, *p* < .001), but not with arousal (β = – 0.19 ± 0.16, t = –1.15, *p* = .25). Figure 3, above, shows the fixed-effect scatter plot, colour-coded by sound clip.

**Figure 3.**
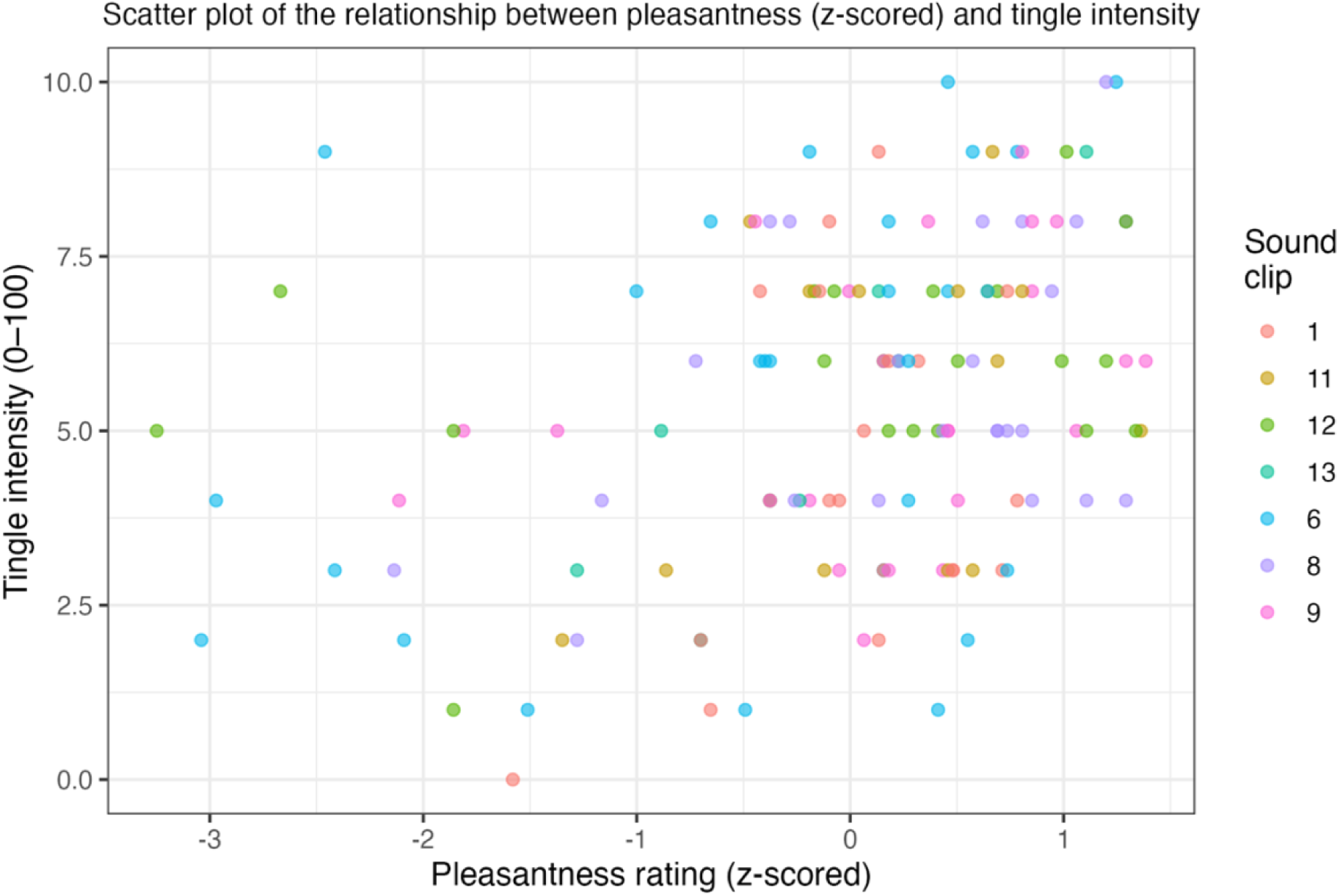
Scatter plot of the relationship between pleasantness (z-scored) and tingle intensity across 136 ASMR-positive trials. Each point represents one trial; colours indicate different sound clips.

A follow-up interaction model revealed that pleasantness remained a strong positive predictor (β = 1.22 ± 0.23, t = 5.25, *p* < .001), while arousal was a significant negative predictor (β = –0.48 ± 0.20, t = –2.39, *p* = .018), and the Pleasantness × Arousal interaction also reached significance (β = 0.40 ± 0.18, t = 2.22, *p* = .028; see Table 2 below).

**Table 2.**
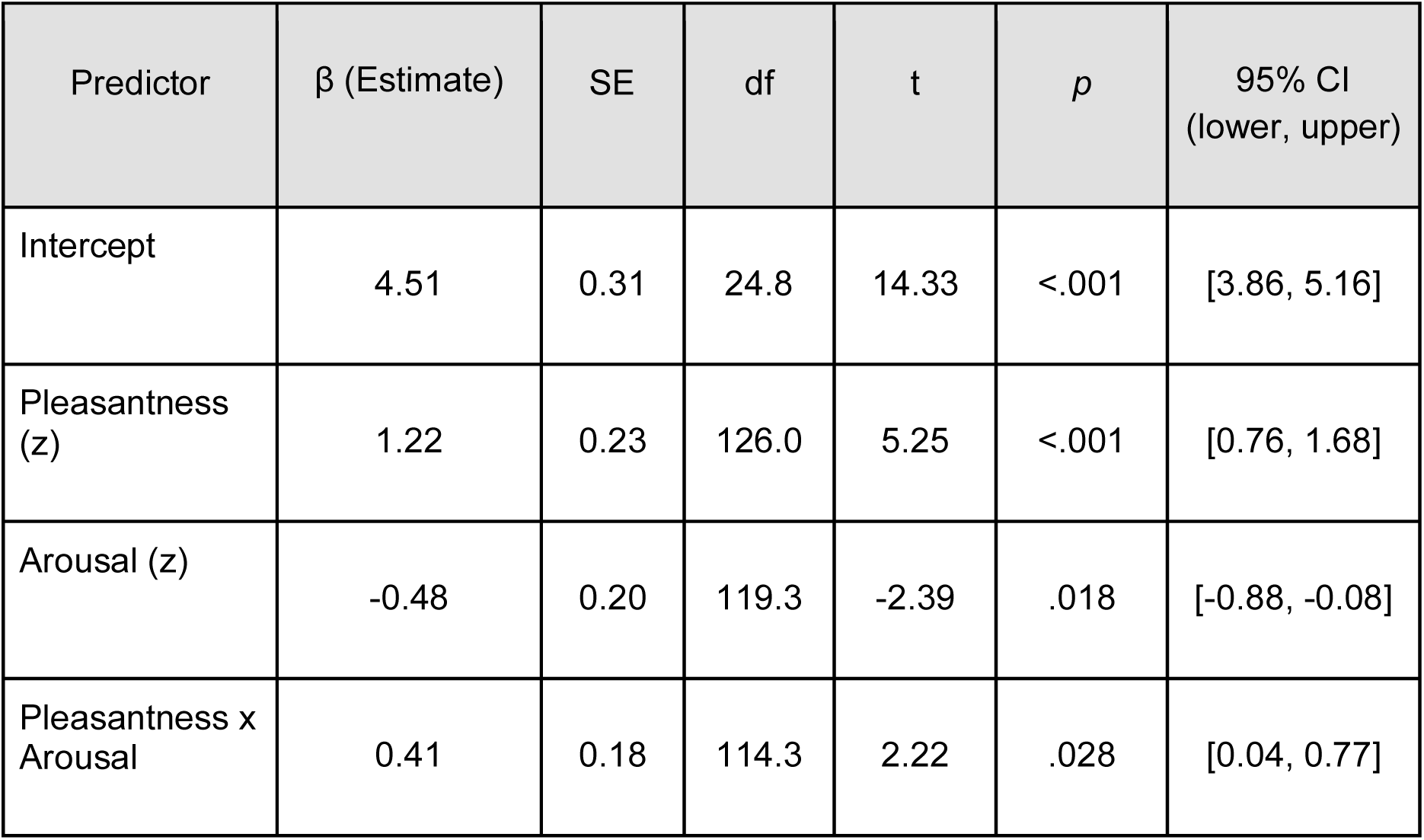
Linear mixed-effects model of tingle intensity ratings (1-10 scale) on ASMR labelled trials. Predictors were z-scored pleasantness and arousal ratings from the affect grid, and their interaction. Random intercepts were included for participant and sound. Pleasantness was a strong positive predictor of tingle intensity, arousal showed a weak negative association, and the Pleasantness × Arousal interaction was significant.

| Predictor | $\beta$ (Estimate) | SE | df | t | p | 95% CI<br>(lower, upper) |
| --- | --- | --- | --- | --- | --- | --- |
| Intercept | 4.51 | 0.31 | 24.8 | 14.33 | <.001 | [3.86, 5.16] |
| Pleasantness<br>(z) | 1.22 | 0.23 | 126.0 | 5.25 | <.001 | [0.76, 1.68] |
| Arousal (z) | -0.48 | 0.20 | 119.3 | -2.39 | .018 | [-0.88, -0.08] |
| Pleasantness x<br>Arousal | 0.41 | 0.18 | 114.3 | 2.22 | .028 | [0.04, 0.77] |

This interaction suggests that the relationship between pleasantness and tingle strength became steeper when arousal was high. In other words, at higher arousal levels, pleasant sounds were more likely to elicit stronger tingles. Conversely, at low arousal, even pleasant sounds were less effective in producing high-intensity ASMR experiences.

### 4.3 Discussion in relation to the Proximity Prediction Hypothesis (PPH)

These behavioural data align closely with key predictions of the PPH, which views ASMR as a vagal cascade triggered by a predicted social-touch event.

#### Valence dominance

ASMR moments are defined by a large hedonic boost and only a minor arousal increase, this dissociation is exactly what would be expected if a slow-stroking CT-touch prediction drives the cascade while sympathetic output is actively suppressed, as the PPH predicts.

#### ASMR experience

The logistic mixed-effects model revealed that pleasantness significantly predicted whether a trial was classified as ASMR, whereas arousal did not. The interaction between pleasantness and arousal was not significant, and model fit was not improved by its inclusion. These findings suggest that the ASMR classification decision relies primarily on the perceived reward value of the sound, a direct prediction of the PPH, and is largely unaffected by concurrent arousal levels.

#### Intensity gradient

Within trials when ASMR was reportedly experienced in response to the sound, the intensity of tingles increased with pleasantness. Furthermore, an interaction emerged where pleasantness was an even stronger predictor of tingle strength when arousal was high.

Under the PPH, this fits the notion that tingle intensity is a graded posterior-insula simulation of predicted touch value, with arousal acting as a gain control mechanism. That is, when arousal is elevated, the system may amplify the hedonic signal, but only when that signal is already strong. These data align with recent behavioural evidence from Gillmeister et al. (2022), who found that tingle intensity during ASMR closely tracked pleasantness and was further amplified by interpersonal touch. This could suggest that affective valence is central to the ASMR simulation, while arousal may act as a gain control mechanism, steepening the link between high pleasantness cues and tingling under certain conditions.

## Conclusion

Taken together, the data suggest that hedonic valence is the primary driver of both ASMR occurrence and intensity. Arousal shows a more equivocal role, sometimes enhancing the pleasantness-tingle gradient, but otherwise exerting weak or inconsistent effects. This ambiguity fits with the PPH view that arousal may reflect both proximity-based alerting and vagally mediated suppression, depending on the listener and context. The remaining discussion in this paper will cover the clinical applications and future tests of the PPH.

## 5. Integrating PPH with CNS-ANS communication and clinical angles

The Proximity Prediction Hypothesis (PPH) describes how a near-ear sound can initiate a cascade that ends in tingles and calm. The present section places that mechanism within wider brain-body communication processes, details the existing physiological clues that suggest the proposed PPH chain is real, and explains why the same mechanism could become a cheaper, accessible alternative to treatments like vagus nerve stimulation (VNS), particularly valuable for anxious and autistic populations as well as though suffering from sleep issues. VNS refers to the implanted, pulse generator therapy in which electrodes are wrapped around the cervical vagus to deliver periodic electrical bursts, a treatment approved for drug-resistant epilepsy and difficult to treat depression already.

### 5.1 Parallels with transcutaneous auricular vagus nerve stimulation (taVNS) and its clinical benefits

Electrical transcutaneous auricular vagus nerve stimulation (taVNS) is a wearable version of implanted VNS. Instead of placing electrodes on the cervical vagus, two small clip electrodes are positioned on the cymba conchae, this is the upper hollow of the outer ear where the auricular branch of the vagus nerve (ABVN) terminates in the skin. A battery-powered stimulator then delivers painless, low frequency pulses (typically 25 Hz, 200-300 µs) for about 15 minutes. Because the ABVN projects directly to the nucleus tractus solitarius (NTS) in the brainstem, the current accesses central vagal pathways without passing through major muscle or bone tissue, unlike cervical VNS. From the NTS the signal ascends to the LC and parabrachial complex and descends to the dorsal motor nucleus of the vagus (DMV), shifting the LC to DMV balance toward parasympathetic dominance. The immediate physiological signature (heart rate deceleration and a rise in high frequency HRV) has been reported to appear within five minutes of stimulation (Borges et al., 2021) and mirrors the pattern ASMR listeners report during tingles.

We propose that taVNS offers a clinical precedent for what the PPH claims ASMR achieves through sensory prediction. Controlled trials have demonstrated that nightly sessions of taVNS significantly enhance sleep quality, reduce insomnia severity, and increase total sleep duration in individuals with chronic insomnia (Zhang et al., 2024; Wu et al., 2022). Similarly, heart rate deceleration of 3-5 bpm and a 5-8% HF-HRV gain has been found during reported ASMR tingling episodes (Poerio et al., 2018) and around 80% of habitual listeners use ASMR to fall asleep (Barratt and Davis, 2015).

Recent studies have demonstrated that brief taVNS courses translate the vagal tone shift into clinically meaningful anxiety relief. In a double-blind, randomized controlled trial, Ferreira et al. (2024) found that a brief taVNS protocol significantly reduced anxiety symptoms in university students, as measured by the Beck Anxiety Inventory, with effects persisting up to two weeks after stimulation. A recent randomized clinical trial by Zhang et al. (2024) found that 8 weeks of taVNS significantly reduced anxiety and depression symptoms, as measured by the Hamilton Anxiety Scale (HAMA) and Hamilton Depression Scale (HAMD) alongside significantly improved Pittsburgh Sleep Quality Index (PSQI) scores. These studies confirm that taVNS can pivot the LC-DMV axis from sympathetic vigilance toward parasympathetic calm and that standard clinical measures of sleep, anxiety, and depression offer realistic indices of this shift. Eid and colleagues (2022) found that ASMR-experiencers, who began with higher baseline state anxiety, experienced a significant reduction in State-Trait Anxiety Inventory-State subscale (STAI-S) scores after viewing an ASMR video, while non-experiencers did not. The PPH therefore predicts that an equivalently large ASMR induced HF-HRV burst should yield comparable anxiety and sleep gains as taVNS. In this way, both taVNS and ASMR seem to converge on a common LC to DMV pathway, but ASMR does so by delivering a precise sensory prior, that ‘pleasant CT-touch is imminent’, instead of an exogenous current.

The analogy yields a clear, testable prediction: the magnitude of an individual’s ASMR induced HF-HRV burst should correlate with reductions in sleep onset latency, and improvement in sleep quality and duration that night, and with drops in state anxiety levels the next day, just as taVNS dose covaries with clinical improvement. Future work can evaluate this prediction with single night polysomnography and standard anxiety inventories such as STAI-S. Currently, taVNS is being trialled as an intervention for treatment-resistant depression, PTSD and insomnia, yet it requires specialised hardware and clinical monitoring. ASMR could offer a headphone based, low cost, surrogate for taVNS, potentially expanding vagal tone interventions to populations who lack access to medical hardware, potentially providing similar clinical benefits that anyone with headphones could utilise.

### 5.2 Why anxious and autistic listeners might benefit most from ASMR-based interventions

A growing evidence base confirms that listeners do not seek out ASMR videos merely for curiosity or entertainment but because the experience delivers measurable relief from anxiety and sleeplessness, ASMR is therefore ripe with potential clinical applications.

How can the predictive coding mechanism underpinning PPH further hone ASMR’s clinical applications to specific populations? Both anxiety disorders and autism spectrum conditions are thought to be characterised by fundamentally over-precise interoceptive prediction errors (Paulus and Stein, 2006; Pellicano and Burr, 2012). In functional terms the insula ‘cries wolf,’ keeping LC tone elevated and vagal tone low. A parallel Bayesian account proposes that autistic brains under-weight priors and over-weight sensory evidence, forcing even mundane events to register as surprising and arousal-worthy (Pellicano and Burr, 2012; Lawson et al., 2014). Both scenarios keep the insula-LC loop chronically engaged. The PPH offers a mechanistic reason listeners from these populations might gravitate toward ASMR videos; having a higher baseline LC tone suggests larger dynamic range for LC suppression, so when the prediction of safe, gentle CT-touch occurs, the resulting vagal release might be steeper, yielding a bigger HF-HRV boost and a more dramatic feeling of relief. Reported experiencers of ASMR have been shown to score higher on neuroticism and anxiety than non-experiencers, suggesting they may have more to gain from a parasympathetic tilt in general (McErlean and Banissy, 2017). Supporting this, Poerio et al. (2021) demonstrated that ASMR experiencers exhibit heightened sensory sensitivity across multiple modalities, including increased bodily awareness and interoceptive sensitivity. Autistic individuals often display atypical interoception; atypical emotional clarity, alexithymia, and interoceptive confusion (Bonete et al., 2023). Sensory processing in autism is also frequently atypical, with both hyper- and hypo-responsiveness across modalities (Elwin et al., 2013), possibly accompanied by somatosensory amplification. Interestingly, some autistic adults report heightened bodily awareness despite reduced interoceptive accuracy, indicating a mismatch between subjective and objective interoceptive states (Garfinkel and Critchley, 2016). This convergence suggests that ASMR may be especially impactful for individuals with enhanced sensory and emotional responsiveness, possibly due to a greater dynamic range in their autonomic regulation potential.

While ASMR proneness also correlates with trait empathic concern (McErlean and Banissy, 2017), this does not preclude its relevance for autistic individuals, who may differ in ‘cognitive empathy’, i.e. imagining another person’s mental state, but not necessarily ‘affective empathy’, the capacity to emotionally resonate with affiliative or caring cues (Dziobek et al., 2008). These findings map onto the PPH cascade: a powerful, but non-intrusive, prior, silences insular error signals, drops LC tone, and brings the body into a parasympathetic state that many autistic and anxious individuals may otherwise struggle to access. If near-ear audio can normalise the LC-DMV balance in these populations, it may again serve as a low cost alternative to taVNS, especially for children or adults who are needle-averse or have restricted access to neurostimulation clinics.

Evidence from tactile research reinforces the clinical logic. Even a single session of massage, can produce immediate reductions in state anxiety, along with decreases in blood pressure and heart rate (Moyer et al., 2004). Scalp massage specifically, in office workers, significantly reduced cortisol, blood pressure, heart rate, and self-reported stress (Kim et al., 2016), suggesting that even brief, localised tactile input can rapidly shift the autonomic balance toward parasympathetic dominance. Yet CT-touch is not always socially available or desired by people with heightened sensory sensitivities. ASMR supplies a predictive, contact-free analogue that can be self-administered with nothing more than headphones for people in anxious or autistic populations who may be otherwise touch-avoidant.

### 5.3 Refining New CAN biomarkers

The PPH model lends itself to proposing two straightforward improvements to the resting HF-HRV score as a biomarker of CAN health, which dominates the current literature. Firstly, instead of looking at HRV in a long, resting baseline, how it changes from trial to trial could be observed while someone is listening to ASMR. A mixed-effects regression of HF-HRV gain and reported tingle intensity will provide a slope for each person. A steep positive slope should indicate that the person’s vagus nerve immediately answers the brain’s ‘this is pleasant and safe’ signal; a flat slope means it does not. If the PPH model is correct, the individual differences in responsiveness are possibly more informative about anxiety risk or sleep quality for that individual than a single resting HF-HRV snapshot. Furthermore, if the PPH is supported, then combining biomarkers like beta band power decreases in the posterior insula (suggesting the system in PPH is predicting a gentle CT-touch is about to happen), along with HF-HRV gain, would provide a mechanistically coherent biomarker that directly indexes the hypothesised cascade from cortical prediction to autonomic change.

Such a multimodal index could predict who will report feeling less anxious or who might fall asleep faster, more reliably than either brain or heart signal could when taken on its own.

In short, the PPH model suggests that ASMR tingling could be used as a convenient stress-test of the CAN loop across individuals, one that can be quantified in real time and may add to the current diagnostic toolkit for anxiety, insomnia, and related conditions. Further experimental paradigms that could be used to test the PPH model are discussed in the next section.

## 6. Future tests of the PPH model

The PPH makes concrete, falsifiable claims about where in the sensory chain the ASMR cascade begins and how it propagates through the insula-LC-vagus axis. Below, a series of experimental predictions, ranging from psychophysics to source-localised MEG, are outlined, to suggest what results would support these claims in future research.

### 6.1 Distance manipulation predictions

#### Binaural morphing of approach cues

The PPH model suggests that the ASMR cascade is gated by perceived proximity, such that a continuous morphing of binaural cues from far (>1m) to near field (<30 cm) should show a non-linear inflection point in ASMR reports, with tingle likelihood, pleasantness, and vagal markers (e.g., HF-HRV gain) rising sharply as the sound enters the peri-aural space. This would reflect the transition into the brain’s peripersonal comfort zone, aligning with prior PPS boundaries observed in audio-tactile studies (Ferri et al., 2015; Serino et al., 2009). Given that pupil dilation has been observed during ASMR listening, likely reflecting heightened attentional engagement with the sound, the PPH further predicts that a delayed pupil constriction should follow as parasympathetic dominance increases during the latter stages of the response. This later-phase constriction has not yet been empirically tested but would be expected if the LC-vagus balance shifts toward sustained calm. This could be tested using interaural time/level difference manipulations of a typically ASMR-inducing stimulus, such as ‘realistic haircut sounds’ (Schürmann et al., 2006).

#### Disrupting spatial coherence across ears

If spatial proximity is integrated across both ears to determine whether the stimulus is near or far, then presenting conflicting distance cues across ears (e.g., one ear hears a close whisper; the other a far-filtered version) should reduce ASMR responses and vagal activity, relative to conditions with coherent near-field input in both ears. This would support the view that the brain uses spatial coherence as a gating signal for engaging the insula-LC-vagus cascade and disrupting it should reduce the probability of experiencing tingles.

### 6.2 Combining real CT-touch with near-ear audio predictions

Recent findings by Gillmeister et al. (2022) indicate that ASMR responders not only exhibit a higher incidence of mirror-touch synaesthesia but also report greater positive emotional reactions to social touch, especially those with stronger ASMR traits. While this supports the notion that affective touch and ASMR share common hedonic mechanisms, the next step is to test whether these effects reflect underlying prediction-based neural dynamics. If the PPH is correct, combining real CT-touch with auditory cues should produce distinct physiological and neurophysiological signatures that reflect audio-tactile congruence and temporal precision.

#### Audio-tactile congruence

Stroking of the listener’s scalp at CT-optimal velocity (3 cm s⁻¹) while presenting either a near-ear brushing sound (congruent) or an identical sound filtered to far-space (incongruent) should boost posterior-insula β-ERD and HF-HRV if the PPH is to be supported. Whereas incongruence will dilute both markers, because the prediction error becomes more precise when the auditory prior and tactile evidence disagree (Ellingsen et al., 2016).

### 6.3 Expectation modulation and proximity cue predictions

Ellingsen and colleagues (2013) devised an elegant ‘placebo-hedonia’ protocol; an inert nasal spray presented as a ‘pleasure enhancer’, followed by slow brush strokes on the forearm during fMRI. The placebo increased subjective pleasantness ratings by around 25%, with enhanced BOLD activity in S1, S2, and the posterior insula, and elevated functional coupling between the pregenual ACC (pgACC) and periaqueductal gray areas, supporting a top-down prediction-based modulation of somatosensory gain. In predictive coding terms, the positive label increased the precision of the ‘this will feel good’ prior, allowing top-down signals to dominate and turn up the gain on the incoming CT volley.

Building on Ellingsen’s finding that positive expectancy amplifies CT-touch processing, if the PPH is correct in asserting that tingle cascades result from precision-weighted predictions of CT-optimal touch, then positively framing a binaural track (e.g. labelling it as a ‘clinically validated tingle inducer’) should increase posterior-insula β-band desynchronization, enhance vagal tone (HF-HRV), lead to a constriction in tonic pupil diameter, and raise subjective ratings of tingle intensity and pleasantness, provided the track contains proximal, near-earl spatial cues. Furthermore, if spatial proximity is a prerequisite for CT-touch predictions, then far-filtered versions of the same track should fail to elicit ASMR responses, even under positive expectancy conditions. That is, labelling alone will not boost tingles or parasympathetic markers when the sensory input lacks coherent proximity information. This prediction sharply distinguishes the PPH from a purely cognitive account: both sensory proximity and cognitive framing must converge to silence prediction error and initiate ASMR.

### 6.4 Predicted EEG and MEG signatures of the PPH cascade

If the PPH is correct, ASMR should elicit a specific neural-autonomic sequence reflecting affective touch simulation and vagal modulation. Empirically, EEG studies show that ASMR triggers produce increased alpha, gamma, and modulations in sensorimotor rhythms (Fredborg et al., 2021) and reduced theta coupled with elevated beta (Engelbregt et al., 2022), along with immediate pupil dilation during strong ASMR episodes (Pedrini et al., 2021). Building on this, PPH predicts a time-locked cascade in the EEG: an initial beta-band suppression over centroparietal sites reflecting S2/posterior-insula activation for CT touch, followed by a transient gamma enhancement. MEG, with better spatial resolution, should localize this beta-gamma sequence to the posterior insula and OP1/S2, with earlier beta suppression in pSTS marking peripersonal space detection, and elevated beta-band coherence between the posterior insula and pgACC/vmPFC regions during the tingling window (reflecting precision-weighted prediction). Crucially, stronger posterior-insula beta suppression should correlate with larger increases in high frequency heart rate variability (HF-HRV), supporting the proposed insula-LC-DMV coupling underlying vagal gain in ASMR. These neural-autonomic patterns should be absent or markedly reduced in control trials without reported tingles, or when identical stimuli are presented with far-field spatial filtering, providing a decisive test of PPH.

## 7. Conclusion

The Proximity Prediction Hypothesis does more than explain an unusual, pleasant tingling sensation; it places the ASMR phenomenon within the LC-vagus system that modern affective neuroscience regards as influential to various physiological and neurological functions, including emotional regulation, stress responses, and even cognitive abilities.

Near-ear sounds appear capable of fooling the brain, leading to emotional modulation, where a sensory cue suppresses the noradrenergic accelerator (the LC), allowing disinhibition of the vagal brake, and ushers both the brain and body into a restful state. The PPH therefore provides a predictive-coding framework specifying when and how the plausible neural routes proposed by previous structural accounts, such as McGeoch and Rouw’s (2020) cross-activation model, might be engaged, and how the characteristic tingling experience could be generated as a result.

The illustrative data reported here offer behavioural support for this framework. Across trials, ASMR experiences were strongly predicted by hedonic valence (pleasantness), not by physiological arousal, and tingle intensity scaled with pleasantness, in a manner that was modestly amplified by arousal. These patterns are consistent with the PPH account of ASMR as a reward-based simulation of safe affective proximity, rather than a state of heightened energetic activation.

If this mechanism is supported in future work, ASMR videos could evolve from quirky bedtime rituals into evidence-based, widely accessible therapeutic interventions for anxiety reduction and sleep promotion. This is especially salient for populations such as those with autism, where prediction error is chronically elevated and conventional relaxation techniques often fail. Moreover, the proposed neural-autonomic markers of beta suppression in the posterior insula, HF-HRV gain, and pupil constriction, could serve as future biomarkers for personalised treatment selection and efficacy tracking. Each paradigm offered in the future research section isolates a different link in the proposed chain; proximity detection, CT-touch prediction, LC suppression, and vagal release. Convergent success across distance manipulation, expectancy modulation, and longitudinal outcome trials would transform PPH from a heuristic into a mechanistically validated account of ASMR and, by extension, into a blueprint for audio-based vagal therapies in mental health. By explicitly integrating predictive coding principles with the neurophysiology of interoception, PPH also offers a broader contribution to our understanding of how the brain regulates the body in response to socially salient sensory cues.

## Author Contributions

JF and CP contributed to the conception of the theory and the design of the original research experiment from which the illustrative data were drawn. JF collected and analysed these data under the supervision of CP and CM. JF drafted the manuscript; CP provided critical feedback on the theoretical content and structure, and CM on the analysis and reporting of the illustrative data section. All authors approved the final version of the manuscript for submission.

## Data Availability Statement

The illustrative dataset analysed for this article is part of a larger unpublished project and is not publicly available at present. Requests to access these data should be directed to the corresponding author.

